# Endogenous feline leukemia virus siRNA transcription may interfere with exogenous FeLV infection

**DOI:** 10.1101/2021.01.12.426481

**Authors:** Elliott S. Chiu, Sue VandeWoude

## Abstract

Endogenous retroviruses (ERVs) are increasingly recognized for biological impacts on host cell function and susceptibility to infectious agents, particularly in relation to interactions with exogenous retroviral progenitors (XRVs). ERVs can simultaneously promote and restrict XRV infections using different mechanisms that are virus- and host-specific. The majority of endogenous-exogenous retroviral interactions have been evaluated in experimental mouse or chicken systems which are limited in their ability to extend findings to naturally infected outbred animals. Feline leukemia virus (FeLV) has a relatively well-characterized endogenous retrovirus with a coexisting virulent exogenous counterpart and is endemic worldwide in domestic cats. We have previously documented an association between endogenous FeLV LTR copy number and abrogated exogenous FeLV in naturally infected cats and experimental infections in tissue culture. Analyses described here examine limited FeLV replication in experimentally infected peripheral blood mononuclear cells. We further examine NCBI Sequence Read Archive RNA transcripts to evaluate enFeLV transcripts and RNA interference precursors. We find that lymphoid-derived tissues, which are experimentally less permissive to exogenous FeLV infection, transcribe higher levels of enFeLV under basal conditions. Transcription of enFeLV-LTR segments is significantly greater than other enFeLV genes. We documented transcription of a 21-nt miRNA just 3′ to the enFeLV 5′-LTR in the feline miRNAome of all datasets evaluated (n=27). Our findings point to important biological functions of enFeLV transcription linked to solo LTRs distributed within the domestic cat genome, with potential impacts on domestic cat exogenous FeLV susceptibility and pathogenesis.

**Importance:** Endogenous retroviruses (ERVs) are increasingly implicated in host cellular processes and susceptibility to infectious agents, specifically regarding interactions with exogenous retroviral progenitors (XRVs). Exogenous feline leukemia virus (FeLV) and its endogenous counterpart (enFeLV) represent a well characterized, naturally occurring XRV-ERV dyad. We have previously documented an abrogated FeLV infection in both naturally infected cats and experimental fibroblast infections that harbor higher enFeLV proviral loads. Using an *in silico* approach, we provide evidence of miRNA-transcription that are produced in tissues most important for FeLV infection, replication, and transmission. Our findings point to important biological functions of enFeLV transcription linked to solo-LTRs distributed within the feline genome, with potential impacts on domestic cat exogenous FeLV susceptibility and pathogenesis. This body of work provides additional evidence of RNAi as a mechanism of viral interference and is a demonstration of ERV exaptation by the host to defend against related XRVs.

## Introduction

Endogenous retroviruses (ERV) are scattered throughout vertebrate genomes, representing 8% of genomic content, with documented impacts on normal biologic processes (Consortium, 2001; Griffiths, 2001). During early stages of endogenization, ERVs accumulate mutations that often render the newly endogenized virus defunct, protecting hosts from potentially deleterious genetic material (Lober et al., 2018). In addition to accumulating mutations, ERVs can act as retro-transposable elements inserting into novel genomic loci. Because ERVs are initiated by intact retroviruses with palindromic long terminal repeat (LTR) flanking sequences, they can be edited from the genome via homologous recombination and other mechanisms that are incompletely understood. Sometimes this process results in remnant genomic segments in the form of solo LTRs (Boeke and Stoye, 1997; Lober et al., 2018). While usually unable to produce infectious virions, many ERVs are still capable of undergoing transcription and may produce functional viral proteins (Knerr et al., 2004; Li and Karlsson, 2016). ERVs also function to enhance and/or promote transcription of proximal host genes. Following fixation in the genome, consequently, ERVs have been usurped by vertebrate hosts for essential biological processes such as placentation, oncogenesis, immune modulation, and infectious disease progression (Crittenden et al., 1987; Knerr et al., 2004; Umemura et al., 2000; Zeng et al., 2014).

ERVs have also been exapted to participate in anti-viral activities against exogenous homologues. Endogenous mouse mammary tumor virus (MMTV)-encoded superantigen negatively selects against self-reacting T-cells, limiting the ability for certain exogenous MMTV strains to infect those T-cells (Holt et al., 2013). Endogenous jaagsiekte sheep retrovirus (JSRV) produces Gag-like proteins that interfere with the regular trafficking mechanisms of exogenous JSRV, thereby reducing viral budding and maturation (Malfavon-Borja et al., 2015). Likewise, endogenous JSRV inhibits cell entry of JSRV through hyaluronidase-2 receptor interference by saturating and ultimately limiting the number of receptors that are displayed on the cell surface (Spencer et al., 2003).

FeLV endogenization has occurred in Felidae of the *Felis* genus, and has been characterized in the domestic cat (*Felis catus*). Eight to twelve nearly full-length enFeLV genomes are present in each genome, with significantly greater numbers of solo LTR remnants (Chiu and VandeWoude, 2020; Powers et al., 2018; Roca et al., 2005). Full-length enFeLV genomes are 86% similar at the nucleotide level to horizontally transmitted exogenous FeLV (exFeLV) (Chiu et al., 2018). Domestic cat exFeLV infects domestic cats across the globe with an incidence that ranges from 3-18% (Bandecchi et al., 1992; Gleich et al., 2009; Muirden, 2002; Yilmaz and Ilgaz, 2000). exFeLV infection has a variety of clinical outcomes, with approximately 60% of infections resulting in aborted or truncated infection, and the remainder progressing to high levels of viremia resulting in hematologic dyscrasias, cancers, opportunistic infections and death (Hartmann, 2011). In a study of a natural FeLV epizootic in a 65-hybrid domestic cat breeding colony, we demonstrated a correlation between higher enFeLV-LTR copy number and cats with regressive or abortive FeLV clinical outcomes. This finding was in contrast to cats with lower LTR copy number, which developed progressive infection and accumulated virulent enFeLV-exogenous FeLV recombinants (Powers et al., 2018). We experimentally infected domestic cat fibroblasts with FeLV and likewise demonstrated that primary cells from cats with greater enFeLV-LTR copy number were more resistant to FeLV infection and viral replication (Chiu and VandeWoude, 2020). This relationship was not observed when we examined FeLV infection and replication related to enFeLV-*env* gene copy number, representing intact full enFeLV genomes, which were found at considerably lower rate of incorporation than enFeLV-LTR (mean of 11 *env* copies/cell versus 57 LTR copies/cell) (Chiu and VandeWoude, 2020).

Cell culture experiments further illustrated highly significant dose-dependent correlation between enFeLV-LTR copy number and viral antigen production, prompting the hypothesis that enFeLV may directly interfere with exogenous FeLV (13). One potential mechanism for direct interference is transcription of enFeLV small non-coding RNAs that regulate gene expression and viral reproduction by degrading target RNA. siRNA, miRNA, and piRNA result in RNA interference (RNAi) via different mechanisms. siRNA and miRNA activate the ribonuclease DICER which processes siRNA and miRNA and incorporates them into RNA-induced silencing complex (RISC) which targets complementary mRNA for degradation (Pratt and MacRae, 2009). Once incorporated into a RISC complex, ssRNA can find its full (siRNA) or partial (miRNA) complementary mRNA strand and signal it for translational repression, mRNA degradation, or mRNA cleavage. A comprehensive review of siRNA and miRNA can be found in (Lam et al., 2015). piRNAs, on the other hand, are typically longer in length compared to siRNA (21-35 nt), which regulate gene expression and fight viral infection using a different mechanism (PIWI-clade Argonautes versus AGO-clade proteins) (Ozata et al., 2019) have been recently demonstrated to contribute to interruption of XRV processes in a newly endogenizing koala retrovirus (KoRV) (Yu et al., 2019). While RNAi is typically considered a potent antiviral mechanism used by plants and invertebrates, there have been evidence that RNAi is also used in mammalian systems to complement their normal antiviral activities governed by first-line interferon responses (Schuster et al., 2019). siRNA has been used to inhibit influenza RNA transcription in chicken embryos and canine cells (Ge et al., 2003). siRNAs have also been demonstrated to be capable of silencing hepatitis A viral infections in non-human primate and human cells (Kusov et al., 2006). Evidence is mounting that miRNA and siRNAs play a role in both promoting and inhibiting HIV replication (Balasubramaniam et al., 2018). As a result, there has been growing interest in research on RNAi as a mechanism of antiviral restriction in humans and other mammals.

To determine mechanisms underlying ERV-XRV interactions in the FeLV system, we used *in silico* approaches to investigate enFeLV transcripts in domestic cat tissues, evaluate transcript abundance and tissue tropism, and assess nature of small RNAs that may function to suppress exFeLV infection. Further, we assessed susceptibility of domestic cat peripheral blood mononuclear cells (PBMC) to examine exFeLV infection compared to fibroblasts. We conclude that enFeLV is transcriptionally active in healthy domestic cats and document significant basal levels of enFeLV siRNA transcription that is tissue specific. enFeLV mRNA and siRNA transcription levels were significantly higher in PBMC in other cells, and we noted significant exFeLV replication restriction in primary PBMC compared to fibroblasts. Our findings provide evidence that enFeLV-LTRs are likely to exert control of FeLV replication via an RNA interference mechanism. We also identified ERV transcripts in domestic cats as well as bobcat (*Lynx rufus*) and Siberian tiger (*Panthera tigris*), indicting transcription of this locus may be linked to an ancient pan-felid retroviral *pol* remnant with anti-retroviral or other functions.

## Results

### PBMCs are less permissive to FeLV infection than fibroblasts

Domestic cat PBMCs derived from six cats and challenged with FeLV attained much lower proviral load levels in culture than domestic cat fibroblast infections (Mann-Whitney U test, p=0.0022; Fig. 2A). At day 5 post-inoculation, the mean proviral load achieved was 7,346 proviral copies of FeLV per million PBMC (range = 958-19,901 proviral copies/million) and only two samples exceeded OD thresholds for positive antigen detection (Fig. 2B). In comparison, fibroblast infections yielded a mean of 262,263 (range = 19,376-1,851,261) proviral copy numbers/million cells on day 5 (Fig. 2A), and CrFK infections resulted in high levels of antigen production compared to PBMC (Fig. 2B).

**Figure 1.**
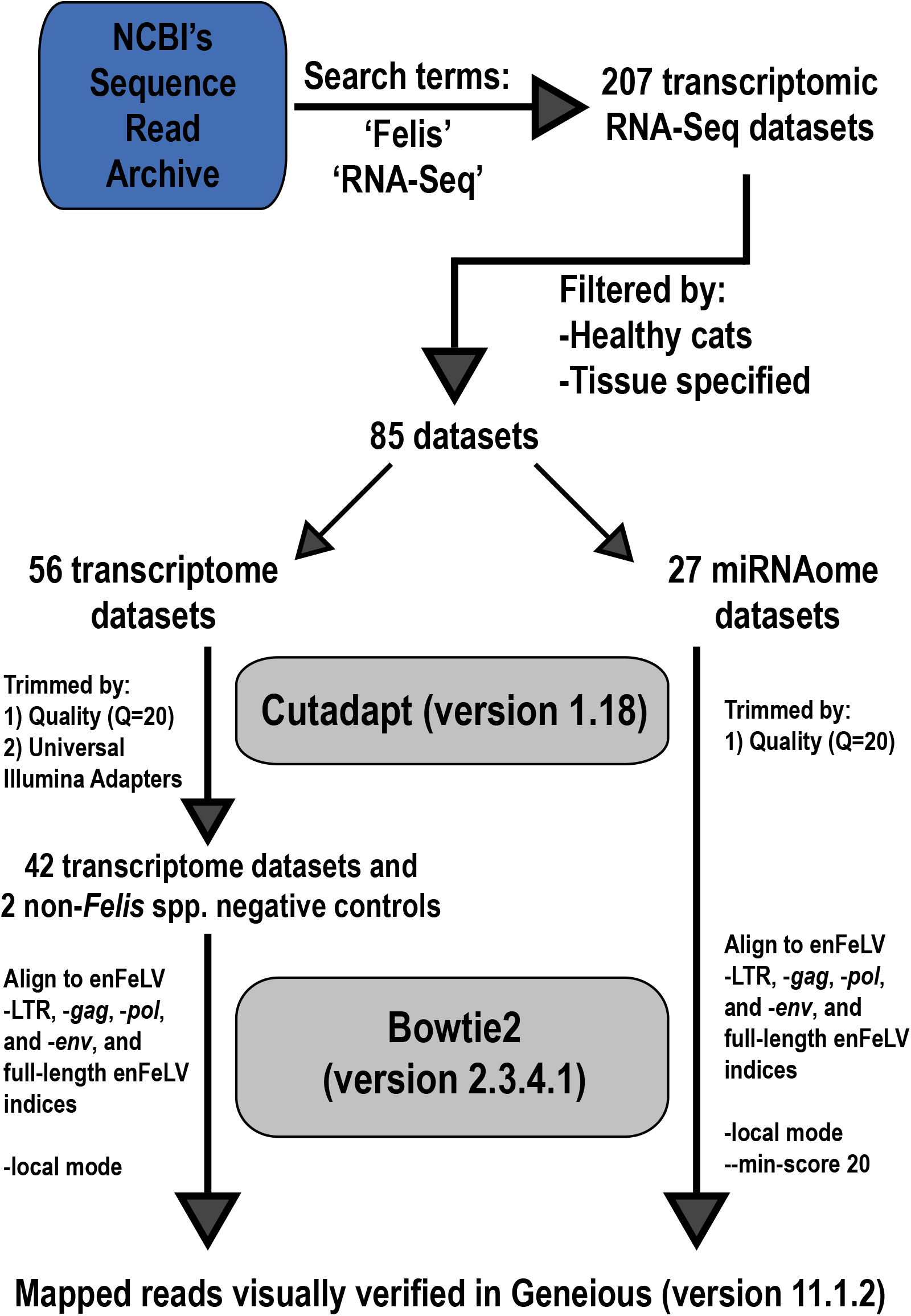
Bioinformatics pipeline used in this analysis. We identified 207 RNA-seq datasets during initial searches. Filtering for healthy cats with defined tissue of origin resulted in 56 datasets for our transcriptome analysis and 27 datasets for our miRNAome analysis. Out of 56 transcriptome datasets, only 33 satisfied quality controls allowing final analysis. Two non-*Felis* spp. transcriptome datasets were included as negative controls.

**Figure 2.**
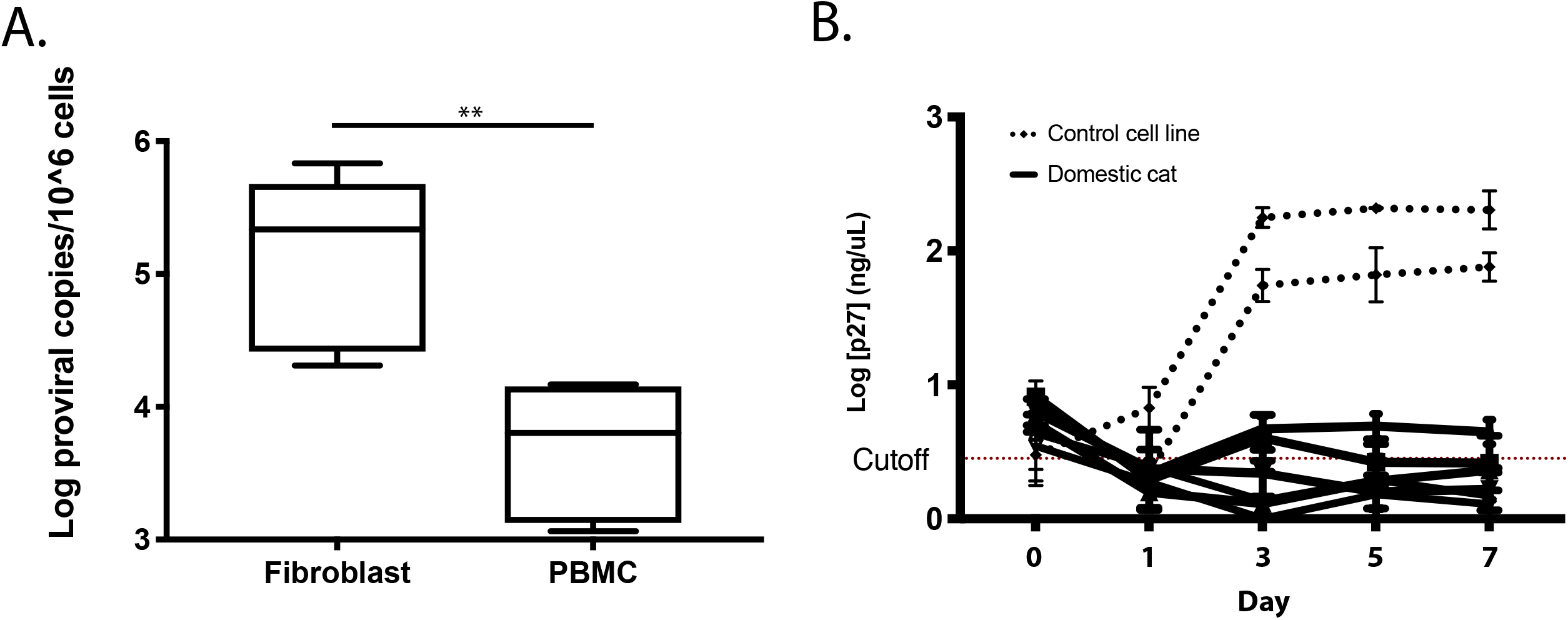
Domestic cat PBMCs were more resistant to FeLV infection than fibroblasts. A) At day 5, median proviral load was 6,359 copies per 10^6^ cells in PBMCS, compared to 119,155 copies per 10^6^ cells in fibroblasts (Mann-Whitney U test; **=p<0.01). B) Domestic cat PBMCs supported low levels of virus replication measured by p27 antigen ELISA. Only two PBMC cultures had transient infections that peaked above the negative cutoff value, established at 3x standard error above average value for negative control replicates (red line). CrFK infections (dotted) were performed as a positive control.

### SRA accessed transcriptome data indicates tissue-specific enFeLV transcription and dominance of LTR transcription

A total of 207 individual animal transcriptomic RNA-Seq datasets were retrieved from the SRA dataset inquiry. Fifty-six of these datasets were from healthy domestic cat tissues of various origins (e.g., embryonic, lymphoid, neural, etc.). Forty-two datasets were included following quality control analysis, representing two studies (99 Lives Cat Genome Sequencing Initiative, *unpublished*) (Fig. 1)(Fushan et al., 2015). An RNA-seq datasets originating from a jaguar (*Panthera onca*) and one from a bobcat (*Lynx rufus*) were used as negative controls (Table S1).

enFeLV transcript levels were approximately 100 reads normalized per million reads (RPM) for most tissues. Outliers included: (1) embryonic tissues (*i.e.* head, body, whole), (2) lymphoid tissues, and (3) a single salivary gland sample. All of these tissues had consistently greater enFeLV transcription levels than neural, skin, reproductive, urinary, lung, digestive, circulatory, and liver tissues (Fig. 3A). Lymphoid (n=4) and salivary gland (n=1) tissues had the greatest enFeLV transcription, averaging approximately 10-fold greater transcription than other tissues.

**Figure 3.**
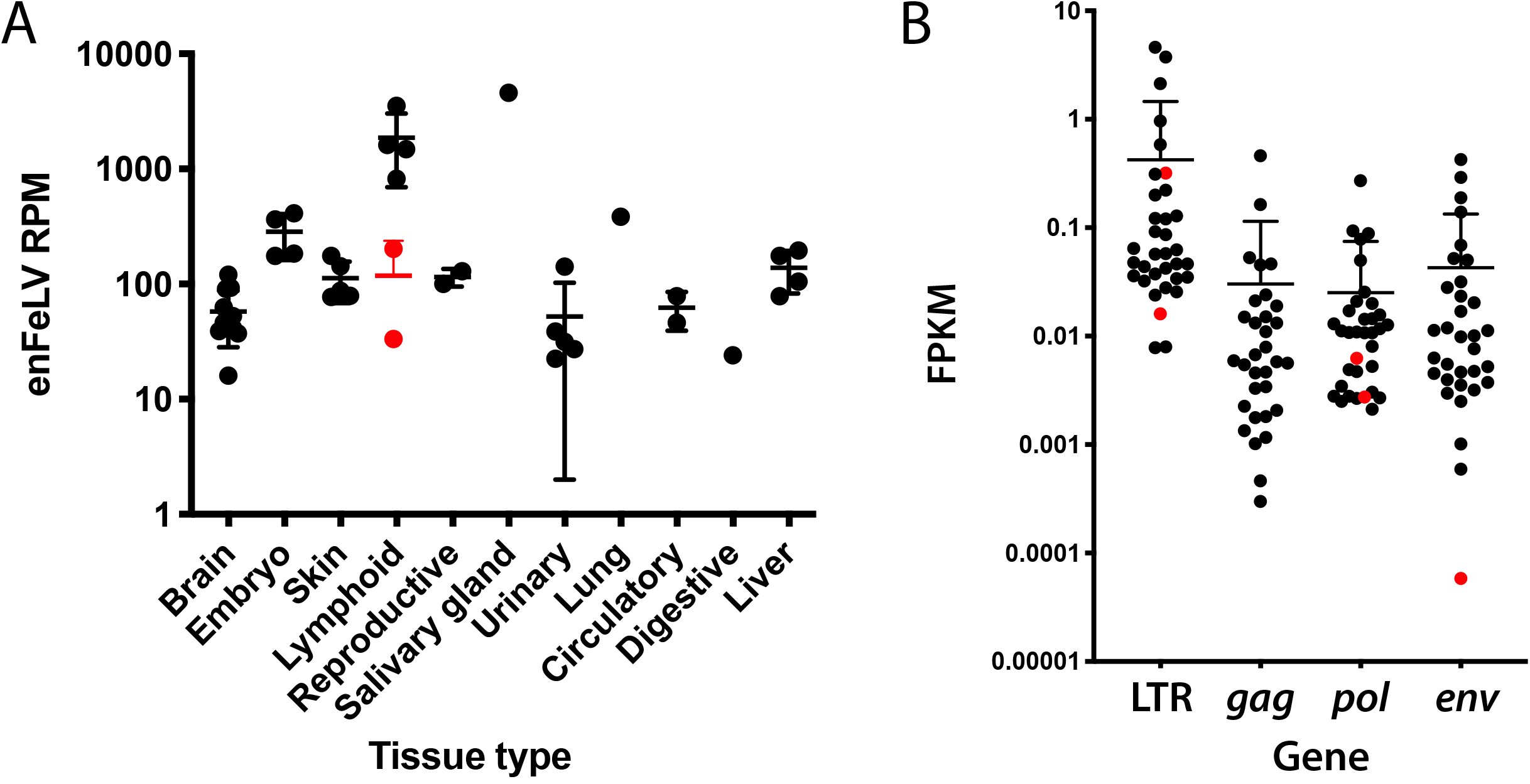
enFeLV transcription is tissues and gene specific. A) enFeLV reads are transcribed at greatest levels in lymphoid and salivary gland tissues. Reads per million (RPM) is a measure of comparison to all other available transcripts in the transcriptome dataset. B) enFeLV-LTR is transcribed at greater levels than other enFeLV genes. Multiple comparisons following ANOVA demonstrated an average 10-fold increase in enFeLV-LTR compared to *gag* (p=0.0079), *pol* (p=0.0073), and *env* (p=0.0111). FPKM = fragments per kilobase million, a measure of total RNA normalized by gene fragment length. Data shown here represent accession numbers SRX211594-211596; SRX211644-211646; SRX211688-211690; SRX1610301-1610326; SRX1625943-1625949. Red data points represent negative control datasets (*P. tigris altaica* – SRX317246; *L. rufus* – SRR6384483).

Following normalization against gene fragment length, enFeLV gene segments were found to have differential expression profiles (Fig. 3B). LTR, *gag*, *pol*, and *env* transcripts represented 0.439, 0.0302, 0.0265, and 0.0453 FPKM of total transcripts, respectively. Relative expression of size-normalized gene segments as FPKM by tissue supports trends identified for full enFeLV; i.e. lymphoid tissues and salivary gland account for the greatest level of transcription (Fig. 4), and LTR transcription is approximately 10-fold greater than other enFeLV genes (Fig. 4A).

**Figure 4.**
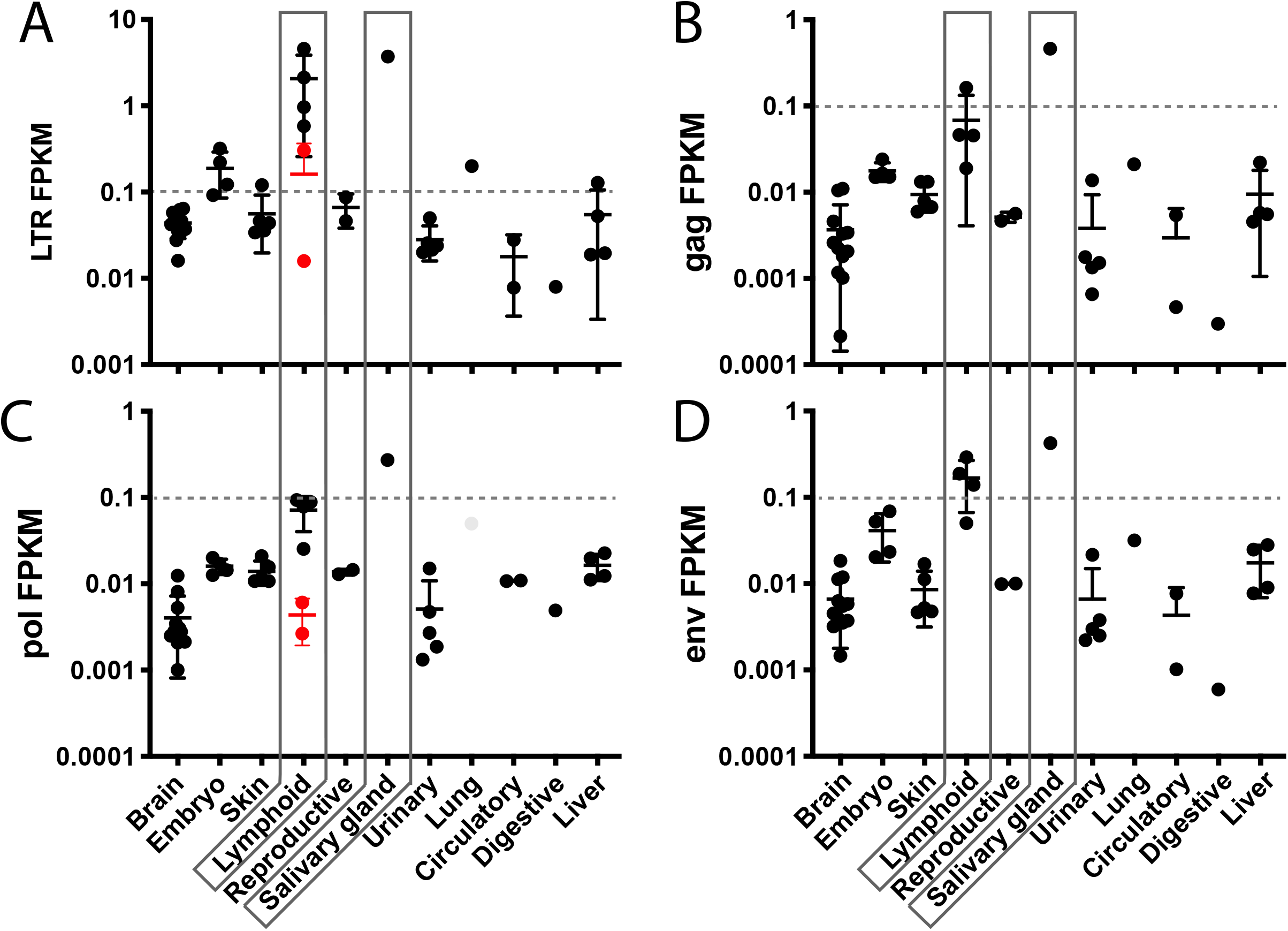
enFeLV genome elements are transcribed variably between all tissue types. A) enFeLV-LTR is transcribed 10 times greater than *gag* (B), *pol* (C), and *env* (D). Lymphoid tissues and the salivary gland (boxed) harbor the greatest amount of enFeLV transcripts across all genome elements. Grey dotted line at 0.1 FPKM are provided for ease of interpreting difference in scale. Data shown here represent accession numbers SRX211594-211596; SRX211644-211646; SRX211688-211690; SRX1610301-1610326; SRX1625943-1625949. Red data points represent negative control datasets (*P. tigris altaica* – SRX317246; *L. rufus* – SRR6384483).

### enFeLV-like RNAs detected in multiple species represent a distinct ERV of felids

RNA datasets from bobcat and Siberian tiger were analyzed as negative controls as these species do not have enFeLV present in their genomes (Polani et al., 2010). We identified two regions that mapped to the enFeLV genome (Fig. 5). One region was found in both bobcat and tiger with short RNA matches driven by a 29-nt poly-adenine stretch in the enFeLV 5′ LTR. The second region varied between bobcat and tiger but both transcripts mapped to an 87-nt region in enFeLV *pol* in the endonuclease/integrase segment of the genome. The tiger transcript was 187-nt and contained 43 SNPs relative to the 87-nt corresponding region of enFeLV *pol*. The bobcat sequence was 179-nt long and contained 39 SNPs relative to enFeLV *pol*. Interestingly, an identical 87-nt region with 100% identity was noted in uncharacterized *Panthera pardus* (Leopard) LOC10927796 mRNA (Accession number: XM_019467717) which was previously reported to represent an ERV (Wei et al., 2011). NCBI’s BLAST tblastn function revealed a shared polymerase gene identity from this region of enFeLV *pol* to feline endogenous retrovirus gamma4-A1 (Accession number: LC176795). Nucleotide similarity between a 90nt region of LC176795 and enFeLV was 60% with 36 SNPs. Pairwise identity between full-length enFeLV (AY364319) and feline endogenous retrovirus LC176795 was 49%.

**Figure 5.**
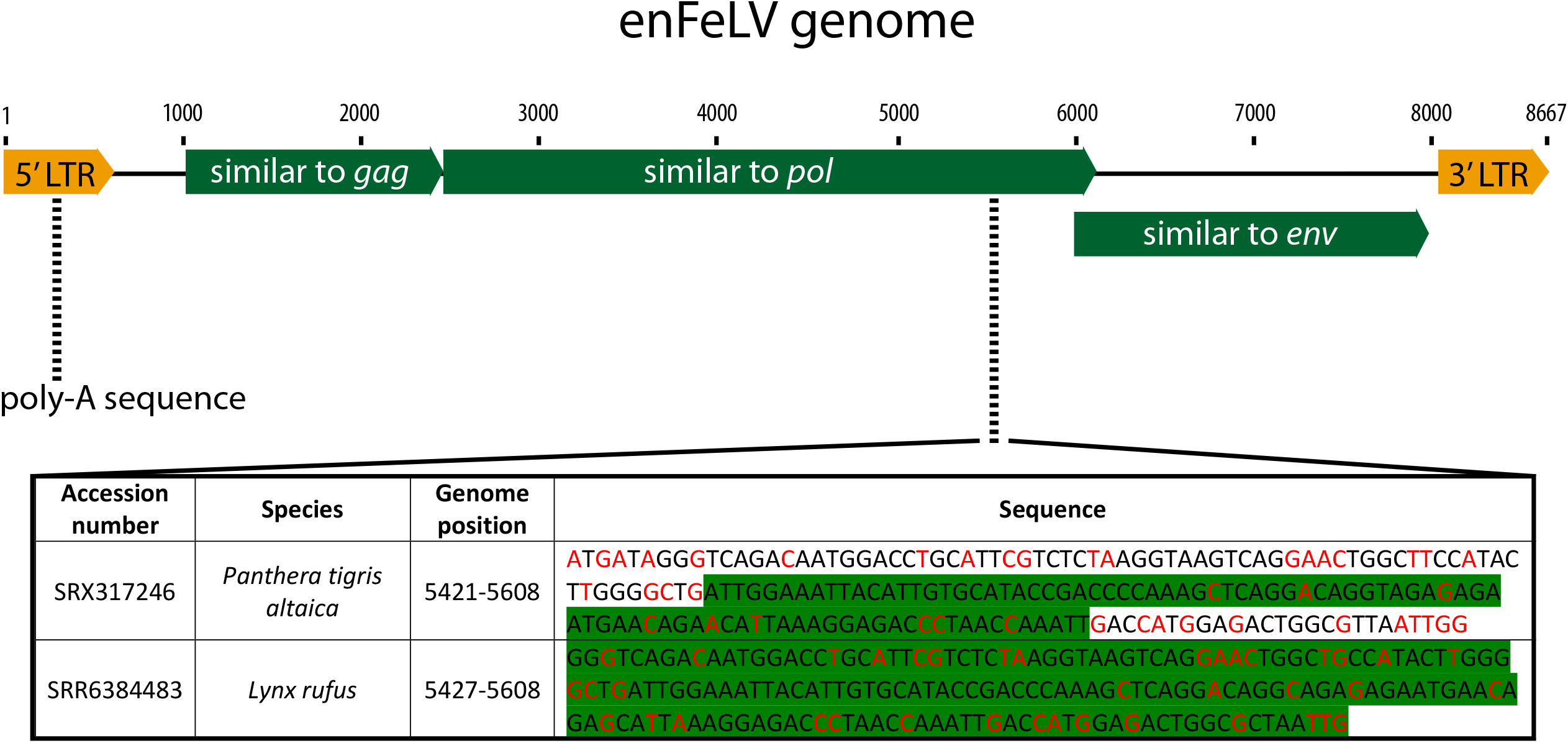
enFeLV genome elements are found in bobcat (SRA Accession number SRR6384483) and Siberian tiger (SRA Accession number SRX317246) transcriptomes. RNA mapped to a poly-A region of the LTR (nt 275-304) and a variable region in *pol* (nt 5,421-5,608). The poly-A region only skewed negative control datasets and did not have an impact on *Felis catus* transcriptome analysis following visual verification. The *pol* mapped reads represented a conserved region that appears to map to an uncharacterized feline endogenous retrovirus that may be distantly related to enFeLV and is found in both the bobcat and Siberian tiger. The sequences highlighted in green were responsible for driving alignment to enFeLV *pol*. Nucleotides highlighted in red represent SNPs.

### SRA accessed miRNAome data identifies abundant enFeLV-derived siRNA transcripts that are both positive and negative sense

Twenty-seven datasets were used to characterize the feline miRNAome from individual animal transcriptomic RNA-Seq datasets retrieved from the SRA (Table S2). These consisted of RNA fragments <30nt and corresponding to RNAs considered to function as RNA silencing transcripts (Lagana et al., 2017). enFeLV miRNA sequences accounted for 0.0163% of all miRNA in the annotated SRA pool. Approximately 75% of the miRNA sequences mapping to enFeLV originate from the LTR (75.1% ± 21.0%), though *gag*, *pol*, and *env* represented more than 10% of the enFeLV-mapped reads in 3-6 of the 27 datasets (Fig. 6, Fig. 7A).

**Figure 6.**
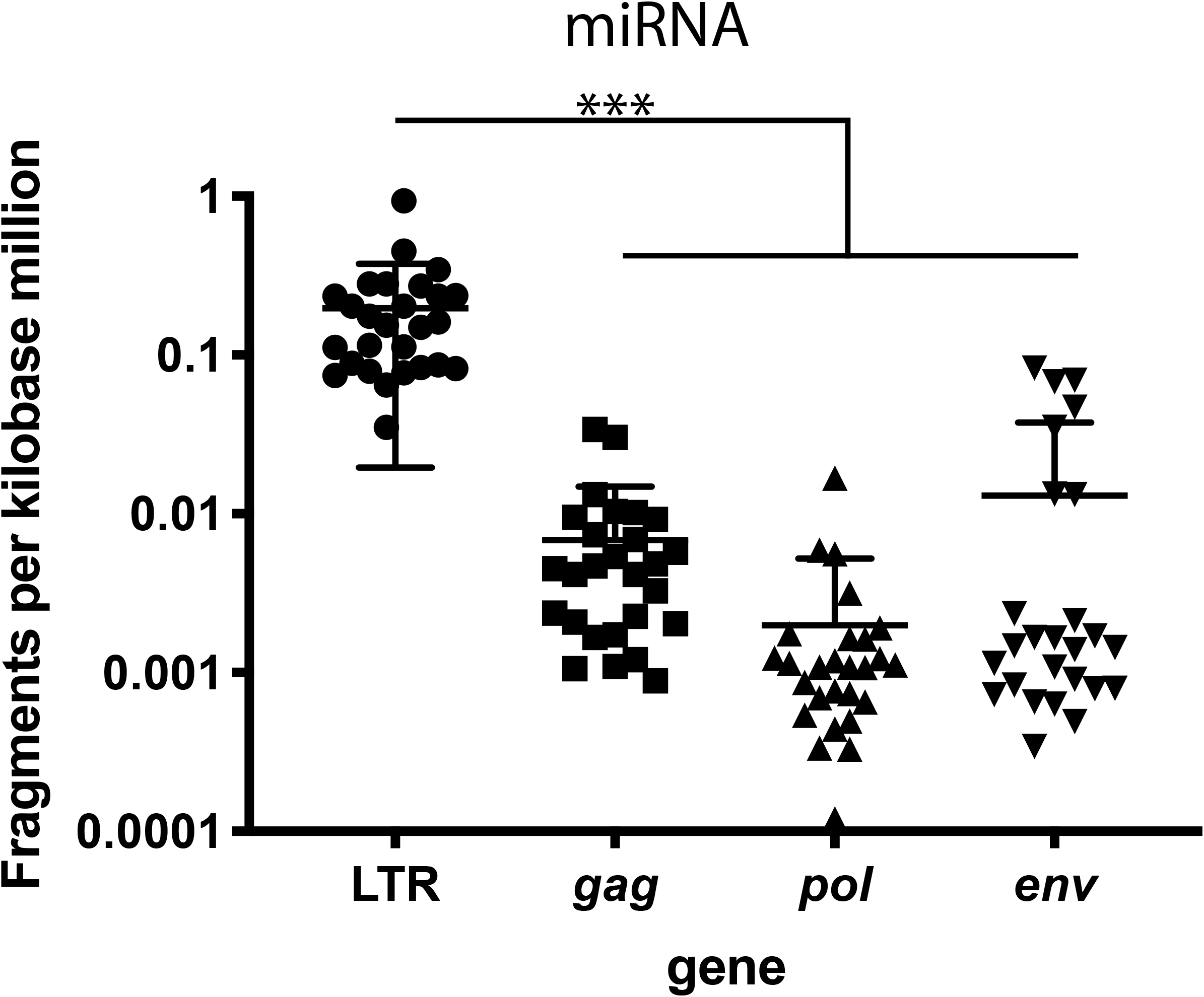
miRNA could be detected for all gene regions in enFeLV but mapped most frequently to enFeLV-LTR (ANOVA; p<0.001). The contribution of miRNA attributed to *gag*, *pol,* or *env* rarely exceeded 0.01 fragments per kilobase million (FPKM). Relative expression was not different among the three genes with background expression proportional to the size of the gene. Increased LTR expression may be driven by both the increased number of LTRs that exist within the genome relative to the other genes and the increased activity of the LTR. Data shown here represent accession numbers SRR4243109-4243135.

**Figure 7.**
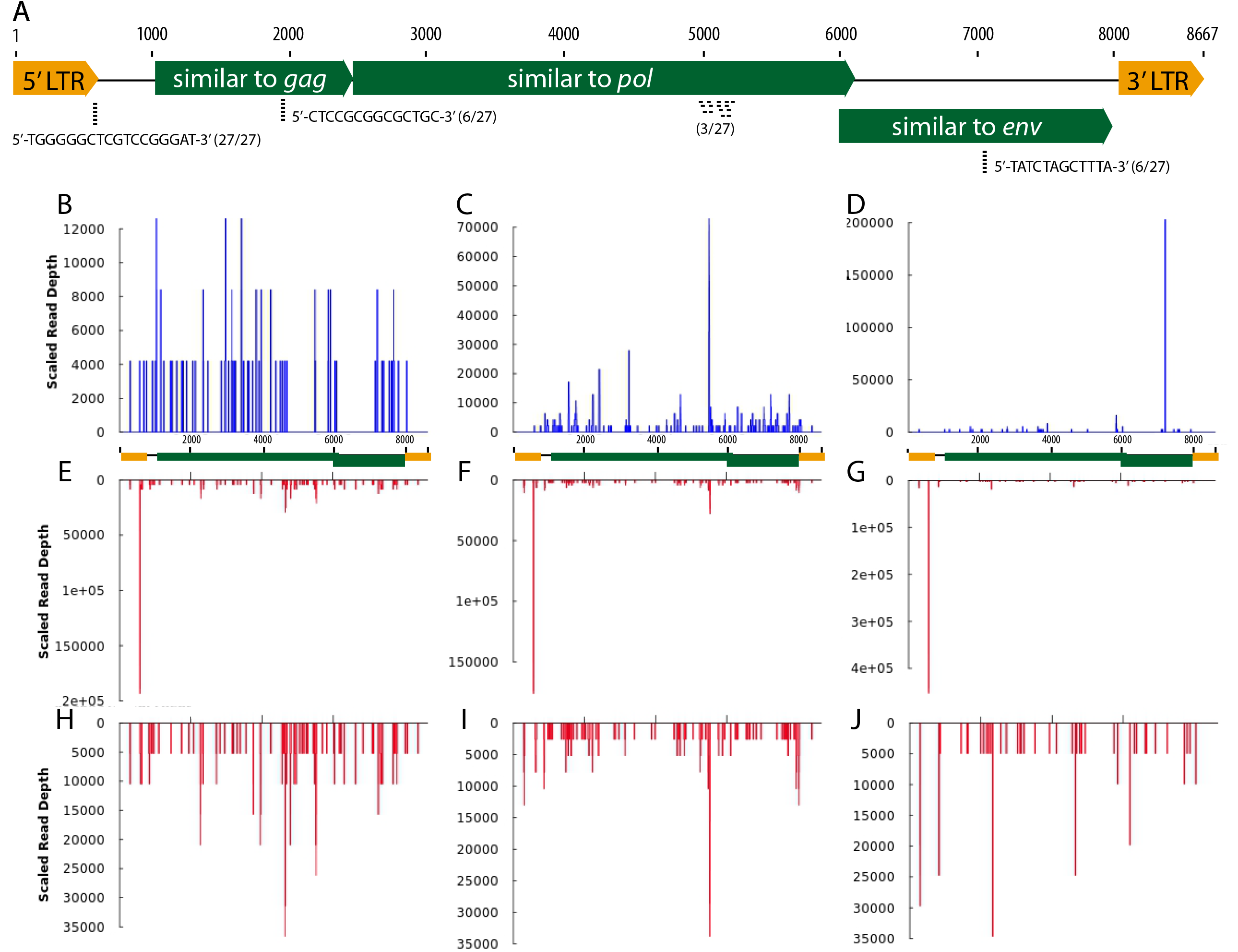
enFeLV miRNA maps to four regions of the enFeLV genome. A) Twenty-seven unique miRNA datasets were evaluated for reads mapping to LTR, *gag*, *pol*, and *env* genome segments. Segments that represented greater than 10% of total mapped reads occurred in 27, 6, 3, and 6 datasets, respectively. Locations of these transcripts are indicated below the genome map, and sequences for these transcripts in LTR (identified in all 27 datasets), *gag* (identified in 6 of 27 datasets) and *env* (identified in 6 of 27 datasets) are indicated. Multiple heterogenous miRNA were identified in 3 of 27 datasets, perhaps indicating contributions from various closely related endogenous retroviruses. B-J) Positive and negative miRNAs identified in three representative datasets are displayed to illustrate individual cat variation in enFeLV miRNA distribution and polarity. Accession number SRR4243132 is represented in Panels B, E, H; Accession number SRR4243126 is represented in Panels C, F, I; Accession number SRR4243130 is represented in Panels D, G, J. Top row (panels B, C, D) indicates positive strand transcript and scaled reads across the enFeLV genome [schematic below the top row indicates enFeLV genome map correlating to map shown in (A)]. Difference in Y axis illustrates variation in scaled read depth from one dataset to another. Panel D illustrates a high abundance transcript that maps to enFeLV *env* in one dataset. Middle row (panels E, F, G) indicates scaled read depth of negative strand miRNA transcripts. The overwhelming abundance of a 21-nt LTR transcript (2-5×10^5^ scaled reads/dataset) obscures lower abundance negative strand transcripts. Removal of this transcript from panels H, I, J allows visualization of lower abundance negative strand miRNAs (5-35,000 scaled reads/dataset) that map to enFeLV.

Characterization of abundant miRNAs: A 21-nt LTR negative-sense miRNA at nucleotide 557 was detected in all 27 individuals and by far the most abundant enFeLV miRNA identified. This transcript is located just 3′ to the 5′-LTR, 74 nucleotides downstream of the transcription start site (Fig. 7A). Sequence for this LTR miRNA, (5′-ATCCCGGACGAGCCCCCACGC-3′), is identical to enFeLV in the same location, with the exception of the 3 flanking nucleotides, and represents a purely negative-stranded population (Fig. 7B,E,H; Fig. S1). The 3 mismatched nucleotides have the lowest sequencing quality score, indicating these are potentially miscalled bases. A 12-nt miRNA segment (5′-TATCTAGCTTA-3′) was identified in *env* at nucleotide 7,045 in 6 of 22 individuals (Fig. 7A). This sequence is positive-stranded and correlates with the gp70 surface protein-like portion of enFeLV Env. Six individuals also had a 14-nt miRNA (5′-CTCCGCGGCGCTGC-3′) at nucleotide 1,963 within the virion core peptide p27 portion of enFeLV *gag* (Fig. 7A). Three individuals also had miRNA segments that mapped diffusely around the endonuclease/integrase region of *pol* (nucleotide region 4800-5200) (Fig. 7A)

sRNApipe miRNA strand polarity analysis was used to analyze strand specificity for miRNA transcription for both abundant and rare miRNA transcripts that exceeded 18-nt in length. Many low copy number miRNA transcripts were found with homology to enFeLV in addition to the primary transcripts noted above. These were generally dispersed across the enFeLV genome. Three representative datasets (Accession numbers: SRR4243126, SRR4243130, SRR4243132) that illustrate unique miRNA transcripts are depicted in Fig. 7B-J, and remaining maps are included in Fig. S1. As noted above, the abundant 21-nt 5′ LTR transcript found in all individuals was a negative-sense transcript, while reads mapping to other regions of the genome were overwhelmingly positive-stranded (Fig. 7B-D, Fig. S1). A unique 20-nt read that was exclusively positive-sense mapped to enFeLV *env* with one deletion and two SNPs in the intervening sequence. miRNA mapping to *pol* is non-specific to a specific locus and has both positive-sense and negative-sense strands mapping to the area.

## Discussion

Despite decades of study, the mechanisms underlying the distinct outcomes of domestic cat FeLV infection remain elusive. The majority of FeLV-infected cats overcome infection, and vaccination can successfully protect against disease, suggesting an adaptive immune response can be protective (Torres et al., 2005). However, a significant proportion of animals exposed to FeLV are unable to eliminate the infection and ultimately succumb to hematologic dyscrasias, lymphoid tumors, or opportunistic infections (Cotter et al., 1975). FeLV replicates to extraordinarily high titers during progressive infection, and in more than 50% of progressive infections, ERV-EXV recombination occurs, resulting in switch in receptor usage and more progressive disease (Powers et al., 2018). Novel observations reported here are highly suggestive that domestic cat enFeLV functions in part to restrict exFeLV infection and provides an explanation for divergent outcomes of FeLV disease.

*In silico* analysis demonstrates that basal enFeLV transcripts are abundant in tissues from healthy cats, enFeLV is transcribed in a tissue-specific manner, and transcript level varies by gene segment (Fig. 3B and 4). LTR transcription is approximately 10 times higher than *pol*, *gag* and *env* (Fig. 4), which may be reflective of the greater number of LTR elements per genome than other segments (Chiu and VandeWoude, 2020). Lymphoid tissue transcription is 1-2 logs greater transcription than other tissues, and one salivary gland transcriptome available for analysis had higher expression than lymphoid (Fig. 4). miRNA transcripts mapping to enFeLV were also detected, including a 21-nt negative-stranded oligoribonucleotide noted in 27 of 27 miRNA transcriptomes in the SRA database (Fig. 7; Fig. S1). FeLV is lymphotropic and has replication phases in salivary tissue that result in viral transmission following social or antagonistic contact (Willett and Hosie, 2013), though here we show that infection in primary PBMC is highly restricted (Fig. 2). Consequently, enhanced enFeLV transcription and basal miRNA production in these tissues may represent a specific host restriction mechanism.

Conversely, higher basal expression of enFeLV in lymphoid tissues may result in a greater potential for ERV-XRV recombination to occur in these cells following co-packaging of endogenous and exogenous transcripts (Stuhlmann and Berg, 1992). Recombination between enFeLV and exFeLV occurs in the 3′ half of the genome in approximately 50% of progressive infections, and is associated with worse clinical outcomes, presumably relating to changes in viral receptor and cell tropism from THTR-1 to PIT-1 (Chiu et al., 2018).

We identified a sizable number short non-coding miRNA transcripts that map to enFeLV sequences. Unlike enFeLV transcription, enFeLV miRNA was present only at specific loci (Fig. 7; Fig. S1), suggesting a specific miRNA function. Given known function of miRNA to degrade complementary mRNA via RISC complex degradation, it seems feasible that these loci represent evolutionarily selected transcript sites that provide host defense against virulent FeLV disease. All 27 cats evaluated were positive for a 21-nt negative-sense miRNA transcript that mapped 3 nucleotides downstream from the 5′-LTR U3 region. The length of 21-nt is indicative of a transcript that functions as an siRNA via an RNAi mechanism as the DICER complex requires a very specific length of RNA (Denli et al., 2004). Mapping this 21-nt sequence to exogenous FeLV demonstrates that 2 or 3 SNPs occur at the 5′ end, which may impact RNAi functionality. Additional evaluation of this specific miRNA and its role in FeLV infection is warranted to assess the capacity of this miRNA to interfere with FeLV infection.

Therefore, one potential explanation for regressive versus progressive FeLV outcomes is as follows:

1. High levels of lymphoid enFeLV-LTR and miRNA transcription pre-empt FeLV infection of PBMC via RNAi-like mechanisms. If miRNA inhibition persists, regression may occur, concurrent with adaptive immune responses that overcome infection.
2. In individuals with lower basal LTR transcription levels (correlating with lower enFeLV-LTR proviral copy number), siRNA restriction mechanisms may fail, resulting in primary infection of lymphoid cells and progressive infection.
3. In other individuals, infection of non-lymphoid tissues with low basal levels of LTR transcription followed by recombination with enFeLV-*env* transcripts may result in XRV-ERV recombinants with enhanced tropism for PBMC. This could result in secondary PBMC infection that potentially overwhelms RNA restriction (again correlating with lower enFeLV-LTR proviral copy number), resulting in progressive infection.

It is likely that these mechanisms operate in conjunction with other more well-understood innate and adaptive antiviral mechanisms to drive FeLV infection outcomes in natural systems.

Subsets of individual animals had predominately positive-sense miRNA that aligned to sites in other FeLV *gag*, *pol*, and *env* genes (Fig. 7; Fig. S1). While positive-sense RNA can be directly used as templates for transcription, positive-sense small RNA is unlikely to do so. Mapped miRNA reads to *gag* and *env* were highly specific to single loci. The size of these specific miRNA is shorter than the RNA recognized RNAi mechanisms require (*gag* – 14nt; *env* – 12nt) and are thus unlikely to participate in RNA silencing using mechanisms that are currently defined. On the other hand, the negative-sense 21nt LTR RNA is complementary to the FeLV sequence and therefore, may serve as the template for FeLV RNA genomes being produced during infection. Furthermore, the scale of the read coverage at the LTR locus compared to the other small RNAs may indicate its biological importance and give us insight into its relative activity. The diffuse miRNA mapping pattern in *pol* may indicate that this transcript may act to silence a broad range of retroviruses with conserved sequences in this region.

It is possible that *gag*, *pol*, and *env* transcripts measured are translated into proteins that contribute to normal biology and physiology as they do in other ERV systems (Johnson, 2019). In 1994, McDougall et al reported that truncated enFeLV Env may participate in direct receptor interference inhibiting exogenous FeLV infection (McDougall et al., 1994). However, retroviral LTRs are not a protein-encoding regions; rather LTRs harbor both promoter and enhancer regions that induce transcription and read-through fusion transcripts that may be processed into functional proteins or other units (Berry et al., 1988). As such, enFeLV-LTR may potentially drive *cis-* or *trans*-activation of host transcription machinery to encode for anti-viral proteins, a process documented in MuLV that relates to ERV-XRV interference (Sanville et al., 2010). This phenomenon has been documented for specific host genes that propagate and inhibit disease processes depending on integration site, including anti-viral proteins such as APOBEC3C (Löwer, 1999; Sanville et al., 2010). If enFeLV-LTRs integrate near anti-viral restriction factors, it could ostensibly prime cells to be more resistant to viral infection. Solo-LTRs formed following ERV retro-transposition may result in fixation of these loci in sites where transcription provides a survival advantage through positive genetic selective. Examination of LTR integration sites in future studies may prove useful in determining what host genes may be impacted by increased transcription or expression.

We noted the interesting phenomenon of relatively higher enFeLV transcription in embryonic tissues (Figures 3, 4). Embryonic ERV transcription has been documented to participate in many normal biological functions. ERV transcription has been inferred as a possible protection mechanism against embryonic viral infections by stimulating innate immunity mechanisms (Grow et al., 2015). ERV expression has also been shown to be important in placentation through syncytins (a syncytium-forming protein responsible for trophoblast invasion of the uterine wall) (Lavialle et al., 2013). During embryonic development of the thymus, host genes (and by consequence, ERVs) are expressed and recognized by the autoimmune regulator (AIRE) so as not to elicit an autoimmune response against ‘self’ proteins (Crittenden et al., 1987). This may allow XRV evade specific adaptive immune responses. Ultimately, discovery of the higher transcription may signal cooption of enFeLV proteins in biological processes.

Transcripts from tiger and bobcat were identified that aligned to the enFeLV genome. One mRNA location in the LTR mapped to a 30-nt poly-A stretch in the 5′-LTR (Fig. 5). *Felis catus* samples did not contain transcripts mapping to this region. A second 90-nt mRNA mapped to nucleotide 5,392-5,488 of *pol* and shares identity to both enFeLV and another endogenous gammaretroviral element described in domestic cats as feline endogenous retrovirus gamma4-A1 (Kawasaki et al., 2017). This 90-nt *pol* endonuclease/integrase region transcript likely represents an ancient conserved motif of retroviral *pol* from an ERV remnant that arose in Felidae prior to divergence between large and small felids. This transcript may represent a *pol* remnant from an ancient endogenized retrovirus with homology to FeLV. The associated LTR segment from this ERV may drive transcription via promotor or enhancer function (Berry et al., 1988). The fact that no other related enFeLV segments were recovered from tiger and bobcat datasets suggests this fragment represents a highly conserved region of *pol* that is transcriptionally active. Future studies to identify additional segments of this ERV may reveal a more ancient ERV than FeLV that spans most Felid species.

SRA datasets were highly valuable for this analysis, but curation of datasets was noted to be highly variable. Lack of description and quality control required us to discard more than 80% of the 207 datasets initially identified. Further, our inquiry has demonstrated that data from non-traditional animal models that provides highly informative comparative data are sorely lacking from genomic databases. For example, the inquiry “HIV RNA-seq” yields 1,491 results as of June 2020, whereas “FIV RNA-seq” yielded only 8 responses. The capacity of comparative genomics studies would be greatly expanded by encouraging analyses that will complement the human datasets available.

FeLV represents a naturally occurring retroviral infection of an outbred species with very well documented clinical and virological outcomes. Here we present compelling evidence that enFeLV-LTR transcripts in feline lymphoid tissue abrogate exogenous FeLV infections. Additional experiments documenting mechanistic aspects of this system in relation to observed natural disease outcomes represents a significant opportunity to understand function of ERV in mitigating viral infections, and to understand mammalian RNA interference mechanisms, impacts of ERV on host evolution, and LTR enhancer and promoter functions that regulate host innate and adaptive immune responses.

## Materials and Methods

### Peripheral blood mononuclear cell infection

Blood was drawn from specific pathogen free domestic cats housed at Colorado State University (IACUC protocol #16-6390A). Peripheral blood mononuclear cells (PBMCs) were isolated from fresh blood by ficoll-gradient centrifugation. PBMCs were cultured in 20% FBS-supplemented RPMI media supplemented with 100 ng/mL IL-2 (Sigma, USA) and 50 ng/mL concanavalin A (Sigma, USA). Primary PBMC cultures were expanded for 2 passages before being directly infected.

PBMCs were plated at 1×10^6^ cells/mL and infected with an MOI of 0.01 FeLV-61E as was previously described (Chiu and VandeWoude, 2020). Briefly, supernatant was sampled at days 0, 1, 3, 5, and 7 and tested for viral antigen using p27 ELISA, as previously described PBMCs were harvested at day 5 to enumerate cell viability and proviral copy number. PBMC proviral and antigen load were compared to fibroblast infection proviral and antigen load conducted simultaneously and reported previously (Chiu and VandeWoude, 2020). Briefly, primary fibroblasts and control Crandell-Rees feline kidney control cells (CrFK) were plated at a density of 50,000 cells per 2 cm^2^ in a 24-well plate and infected with an MOI of 0.01 FeLV-61E. Supernatant was sampled at days 0, 1, 3, 5, 7, and 10 and cells were harvested at day 5 and 10 to enumerate cell viability and proviral copy number.

### Endogenous FeLV transcriptomic analysis

Domestic cat transcriptome and miRNAome data sets were acquired through the search function in the NCBI Sequence Reads Archive (SRA) using the search key words: “felis” and “rna-seq.” Datasets were included in the study if they were derived from healthy cats, identified the tissue of origin, and represented transcriptome (excluding miRNAome) datasets (Table S1). Two additional non-*Felis spp* felid transcriptome datasets (*Lynx rufus*, *Panthera tigris*; Accession numbers SRR924676 and SRR6384483/ SRR6384484) were included as negative controls that would not be expected to harbor enFeLV transcripts (Table S1). Tissues analyzed included embryonic (fetus, embryo body, embryo head), neural (cerebellum, parietal lobe, occipital lobe, temporal lobe, hippocampus, spinal cord, retina), skin (skin, ear tip, ear cartilage), lymphoid (spleen, lymph node, bone marrow, thymus), and other organ (muscle, liver, uterus, kidney, testes, pancreas, heart, salivary gland) tissues. All data processing and analysis was completed using the Colorado State University College of Veterinary Medicine and Biomedical Science server. Transcriptome datasets were analyzed using a custom bioinformatics pipeline (Fig. 1). Reads were trimmed for appropriate adapters and by quality (q=20) using cutadapt (version 1.18). The first 600-nt and last 600-nt of the full-length enFeLV were discarded prior to creating the index due to the potential for transcripts to map to other host genomic elements that surrounded the enFeLV integration site. Indices were first generated for full-length domestic cat enFeLV (Accession Number: AY364319) and individual enFeLV gene regions, including LTR, *gag*, *pol*, and *env* separately using Bowtie2-build function (version 2.3.4.1). Transcriptome sequences were mapped to indices using “--sensitive” settings in local mode in Bowtie2 to allow for heterogeneity among different enFeLV genotypes. Alignments were visually inspected by importing mapped .*sam* files into Geneious 11.1.2. Exogenous FeLV was ruled out as the source of mapped reads by looking for exogenous FeLV-specific DNA segments. Any transcripts mapped to the negative controls were manually inspected and their identities confirmed using NCBI’s BLAST tblastn function. Transcriptome reads mapping to full-length FeLV were reported as reads per million reads (RPM). While the reads were mapped as paired-end reads, reported RPM was calculated as unpaired reads. Individual genome elements were reported as fragments per kilobase million (FPKM) to normalize for size of the respective gene region (full-length enFeLV-8,448nt; LTR – 592nt; *gag* – 1,512nt; *pol* – 3,630nt; *env* – 2002nt). Percent transcription for full-length FeLV was analyzed by ANOVA in Prism (version 7.0). Custom scripts are available at https://github.com/VandeWoude-Laboratory/Florida-panther-virus.

### Feline miRNAome analysis

miRNAome datasets were accessed as described above and by querying “felis” and “rna-seq” that also identified miRNA-Seq in the library preparation strategy (Fig. 1). Tissues analyzed included neural (cerebellum, cerebral cortex, brain stem), skin (skin, lip, tongue), lymphoid (spleen, lymph node), and various organ (pancreas, kidney, liver, lung, testis, ovary) tissues. Using the full-length enFeLV index described above constructed with Bowtie2-build, miRNAome reads were mapped on local mode with a minimum threshold score set at 20 in Bowtie2 to account for miRNA’s intrinsically short length. miRNA reads mapping to full-length enFeLV were reported as percent reads mapped to each genome element compared to total mapped reads to the full-length FeLV genome. Percent miRNA mapped to enFeLV was analyzed by ANOVA in Prism (version 7.0). Custom scripts are available at https://github.com/VandeWoude-Laboratory/Florida-panther-virus.

miRNAome datasets were visualized in strand-specific orientation using sRNAPipe on the Galaxy platform (R. et al., 2018). Default settings were used though three genome mismatches were allowed to accommodate identified SNPs at the 3′-end of a 21-nt LTR siRNA sequence. miRNA was mapped against the enFeLV genome (AY364319), with LTR signifying the only transmissible element input file, and protein encoding genes as the transcripts and mRNA input files. All 27 datasets were analyzed to determine positive and negative strand miRNAs of a minimum of 18-nt in length. A second separate negative-sense analysis was conducted that excluded a highly transcribed 21-nt LTR siRNA in all datasets so that less abundant transcripts could be readily visualized.

## Acknowledgements

This work was supported by National Science Foundation–Ecology of Infectious Diseases award 1413925 and by the Office of the Director of the National Institutes of Health under awards T32OD012201 and F30OD023386. The content is solely the responsibility of the authors and does not necessarily represent the official views of the National Institutes of Health. The funders had no role in study design, data collection and interpretation, or the decision to submit the work for publication. We would like to thank Matthew Moxcey and Tyler Eike for their IT assistance.

---

**Figure S1. enFeLV miRNA maps to the enFeLV genome in the 27 datasets.** Each column of three represent the data from one miRNAome dataset not displayed in Fig. 7 (Accession numbers SRR4243109-4243125; 4243127-4243129; 4243131; 4243133-35). Top rows (blue; panels A, B, C, J, K, L) indicates positive strand transcript and scaled reads across the enFeLV genome [schematic below the top row indicates enFeLV genome map correlating to map shown in (A)]. Difference in Y axis illustrates variation in scaled read depth from one dataset to another. Middle rows (red; panels D, E, F, M, N, O) indicates scaled read depth of negative strand miRNA transcripts. The overwhelming abundance of a 21-nt LTR transcript (2-5×10^5^ scaled reads/dataset) obscures lower abundance negative strand transcripts. Removal of this transcript from panels G, H, I, P, Q, R allows visualization of lower abundance negative strand miRNAs that map to enFeLV.

